# Glutathione contributes to plant defense against parasitic cyst nematodes

**DOI:** 10.1101/2021.09.23.461554

**Authors:** M. Shamim Hasan, Divykriti Chopra, Anika Damm, Anna Koprivova, Stanislav Kopriva, Andreas J. Meyer, Stefanie Müller-Schüssele, Florian M.W. Grundler, Shahid Siddique

## Abstract

Cyst nematodes (CNs) are an important group of root-infecting sedentary endoparasites that severely damage many crop plants worldwide. An infective CN juvenile enters the host’s roots and migrates towards the vascular cylinder, where it induces the formation of syncytial feeding cells, which nourish the CN throughout its parasitic stages. Here, we examined the role of glutathione (L-γ-glutamyl-L-cysteinylglycine, GSH) in *Arabidopsis thaliana* upon infection with the CN *Heterodera schachtii*. Arabidopsis lines with mutations *pad2, cad2*, or *zir1* in the glutamate–cysteine ligase (*GSH1*) gene, which encodes the first enzyme in the glutathione biosynthetic pathway, displayed enhanced CN susceptibility, but susceptibility was reduced for *rax1*, another GSH1 allele. Biochemical analysis revealed differentially altered thiol levels in these mutants that was independent of nematode infection. All GSH-deficient mutants exhibited impaired activation of defense marker genes as well as genes for biosynthesis of the antimicrobial compound camalexin early in infection. Further analysis revealed a link between glutathione-mediated plant susceptibility to CN infection and the production of camalexin upon nematode infection. These results suggest that GSH levels affects plant susceptibility to CN by fine-tuning the balance between the cellular redox environment and the production of compounds related to defense against infection.

## Introduction

Thiol-containing compounds are central components of many pharmacological and biochemical reactions. In its reduced form, glutathione (L-γ-glutamyl-L-cysteinyl-glycine, GSH) is the most abundant low-molecular-weight thiol in most cells. The thiol group of the central cysteine residue, which has a nucleophilic character, is required for reduction and conjugation reactions in which glutathione plays pivotal roles. Besides maintaining tight control of cellular redox status via its reducing and antioxidant properties, glutathione mediates many other physiological processes such as cellular signaling, thiol-disulfide interchange reactions, and xenobiotic metabolism and serves as a major component of the cysteine pool (Noctor et al., 2012).

Glutathione biosynthesis generally occurs via two ATP-dependent steps (Meister, 1995). In the first reaction, a peptide bond forms between the α-amino group of L-cysteine (L-Cys) and the γ-carboxyl of L-glutamate (L-Glu) via a process catalyzed by glutamate–cysteine ligase (GSH1). In the second step, glutathione synthetase (GSH2; Hell and Bergmann, 1990) adds glycine (Gly) to the dipeptide γ-glutamyl-cysteine (γ-EC), to form GSH. In Arabidopsis (*Arabidopsis thaliana*), these two enzymes are encoded by single-copy genes; GSH1 is exclusively localized to chloroplasts, whereas GSH2 is predominantly located in the cytosol and to a lesser extent in plastids (Wachter et al., 2005, Pasternak et al., 2008). Under stress conditions, reduced glutathione is rapidly converted into the oxidized form glutathione disulfide (GSSG), leading to an imbalance in the glutathione redox potential (*E*_GSH_), which is considered an indicator of oxidative stress. The accumulation of GSSG is counteracted by glutathione reductases, which reduce GSSG back to GSH at the expense of electrons provided by NADPH (Marty et al., 2009, 2019).

Glutathione also functions as a co-substrate in conjugation and detoxification reactions catalyzed by glutathione transferases (GSTs), and in glutathionylation of protein thiols as a post-translational protein modification (Dixon et al., 2005). In addition, glutathione acts in plant perception systems to activate basal defense responses that help counter microbial attack (Glazebrook and Ausubel, 1994; Parisy et al., 2007). In coordination with cysteine, glutathione plays a role in inducing pathogen-associated molecular pattern (PAMP)-triggered immunity (PTI) via the recognition of invariant microbial epitopes (PAMPs) by pattern recognition receptors (PRRs; Jones and Dangl, 2006; Alvarez et al., 2011).

Null mutants of *GSH1* in Arabidopsis are embryo-lethal (Cairns et al., 2006). Less severe mutations within this gene appear in five distinct mutant alleles (*rml1, rax1, pad2, cad2*, and *zir1*) that were identified through forward genetic screens and found to show a partial decrease in glutathione production (Glazebrook and Ausubel, 1994; Cobbett et al., 1998; Ball et al., 2004; Vernoux et al., 2000; Shanmugam et al., 2012; Bangash et al., 2019). Studies of these *GSH1*-deficient mutants have shed light on the diverse roles of glutathione in many cellular processes, including plant development and responses to abiotic and biotic stimuli (Potters et al., 2002; Noctor, 2006; Foyer and Noctor, 2011). The Arabidopsis *root-meristemless1* (*rml1*) mutant possesses <5% of wild-type levels of foliar glutathione and exhibits severe defects in plant development (Vernoux et al., 2000). Another Arabidopsis *GSH1* mutant, *zinc tolerance induced by iron 1* (*zir1*), contains <15% of wild-type levels of glutathione in leaves, leading to deficits in Fe-dependent Zn tolerance and impaired nitric oxide (NO)-mediated Fe-deficiency signaling (Shanmugam et al., 2012; 2015). The *cadmium-sensitive2* (*cad2*) mutant contains ∼30% of wild-type glutathione levels and shows sensitivity to cadmium and enhanced susceptibility to *Phytophthora brassicae* (Howden et al., 1995; Cobbett et al., 1998; Parisy et al., 2007). The *regulator of ascorbate peroxidase2 1* (*rax1*) mutant contains ∼40% of wild-type levels of glutathione in leaves and shows increased sensitivity to light stress (Ball et al., 2004). However, the altered glutathione content in *rax1* does not appear to affect plant resistance to *Pseudomonas syringae* or *P. brassicae* (Parisy et al., 2007). The *phytoalexin deficient2* (*pad2*) mutant, the most extensively studied mutant in an allelic series of *GSH1* mutants, possesses only ∼20% of wild-type levels of glutathione in leaves and shows enhanced susceptibility to many pests and pathogens (Dubreuil-Maurizi and Poinssot, 2012). Despite substantial evidence for the role of glutathione in different plant pathosystems, little is known about how glutathione mediates the molecular dialog during plant–nematode interactions.

Plant-parasitic cyst nematodes (CNs) are among the most damaging plant pests and pathogens, causing substantial yield losses globally (Savary et al. 2019). Infective-stage juveniles (J2) of CNs hatch from eggs upon stimulation by mostly unknown host triggers and migrate toward the roots. The CNs make numerous perforations in the epidermal cell wall via back-and-forth movements with their stylets, and invade the roots of the host plant near the root tip. Subsequently, they migrate intracellularly through the root cortical cells to the vascular cylinder. Upon reaching the vascular cylinder, the nematodes probe single cambial or procambial cells to induce the formation of an initial syncytial cell (ISC) as a feeding site (Sobczak et al., 1999). The nematodes inject a cocktail of secretions into the ISC through their hollow stylets to modify plant morphogenetic pathways towards the development of the feeding site (Wyss and Zunke, 1986; Hewezi and Baum, 2013). Hundreds of adjacent root cells successively fuse with the ISC via local cell wall dissolution to form a hypertrophied, multinucleate, hypermetabolic syncytial nurse cell (Wyss and Grundler, 1992).

The juvenile nematode rapidly becomes sedentary due to cell-specific muscle atrophy and starts feeding on the syncytium, which acts as a nutrient sink throughout its parasitic stages (Han et al., 2018). Syncytium development is accompanied by extensive metabolic, transcriptomic, and proteomic changes in the infected root tissues (Siddique et al., 2009; Szakasits et al., 2009; Hofmann et al., 2010; Hütten et al., 2015). The juveniles feed, enlarge, and molt three times to differentiate into males or females. Female nematodes grow rapidly and burst through the root surface, while males regain a vermiform body shape and mobile form by remodeling their neuromuscular structures, leave the roots, and search for females (Han et al., 2018). The female dies after fertilization, and the body wall tans to form a typical brown cyst that envelops and shields the next generation of eggs. The eggs are able to survive for prolonged periods (up to 20 years) in the soil until a suitable host is found growing nearby (Grainger, 1964).

The beet CN (*Heterodera schachtii*) is a detrimental pest of sugar beet worldwide. *H. schachtii* infects over 200 plants from 20 different families, including the model plant Arabidopsis (Sijmons et al., 1991). Here, using the Arabidopsis and *H. schachtii* model system, we investigated the role of glutathione in orchestrating host defense responses to CN infection. We conclude that glutathione contributes to plant defense against CN infection through modulation of cellular redox homeostasis and camalexin production in host roots.

## Results

### Cyst nematode infection activates GSH biosynthesis genes in Arabidopsis

To investigate the roles of glutathione biosynthetic genes during different phases of Arabidopsis parasitism by CN, we examined the expression patterns of *GSH1* and *GSH2* in publicly available transcriptomic data (Szakasits et al., 2009; Mendy et al., 2017). These surveys revealed a significant increase in *GSH1* expression during the migratory (10 hours post-inoculation, hpi) and sedentary stages (5 and 15 days post-infection, dpi) of cyst nematode infection (**Supplementary Table S1**). In comparison, although *GSH2* transcript abundance increased significantly during the migratory stage, it remained unchanged during the sedentary stage of infection.

To validate these microarray data, we analyzed the expression of GSH biosynthesis genes in Arabidopsis root segments infected with CN. We collected several hundred root segments (∼0.2 cm) containing infection sites at 10 hpi (migratory stage) and syncytia at 10 dpi (sedentary stage), and analyzed *GSH1* and *GSH2* expression in these tissues compared with uninfected wild-type Columbia (Col-0) roots by reverse transcription quantitative polymerase chain reaction (RT-qPCR). The results confirmed that *GSH1* transcript levels increased during both the migratory and sedentary stages of CN infection (**Fig. 1A**), whereas *GSH2* was significantly induced at 10 hpi but not at 10 dpi (**Fig. 1B**). Overall, these results suggest that GSH biosynthesis genes are activated upon CN infection, and that the regulation of *GSH1* is more pronounced than that of *GSH2* during CN infection; therefore, we focused our further analysis on *GSH1*.

**Fig. 1.**
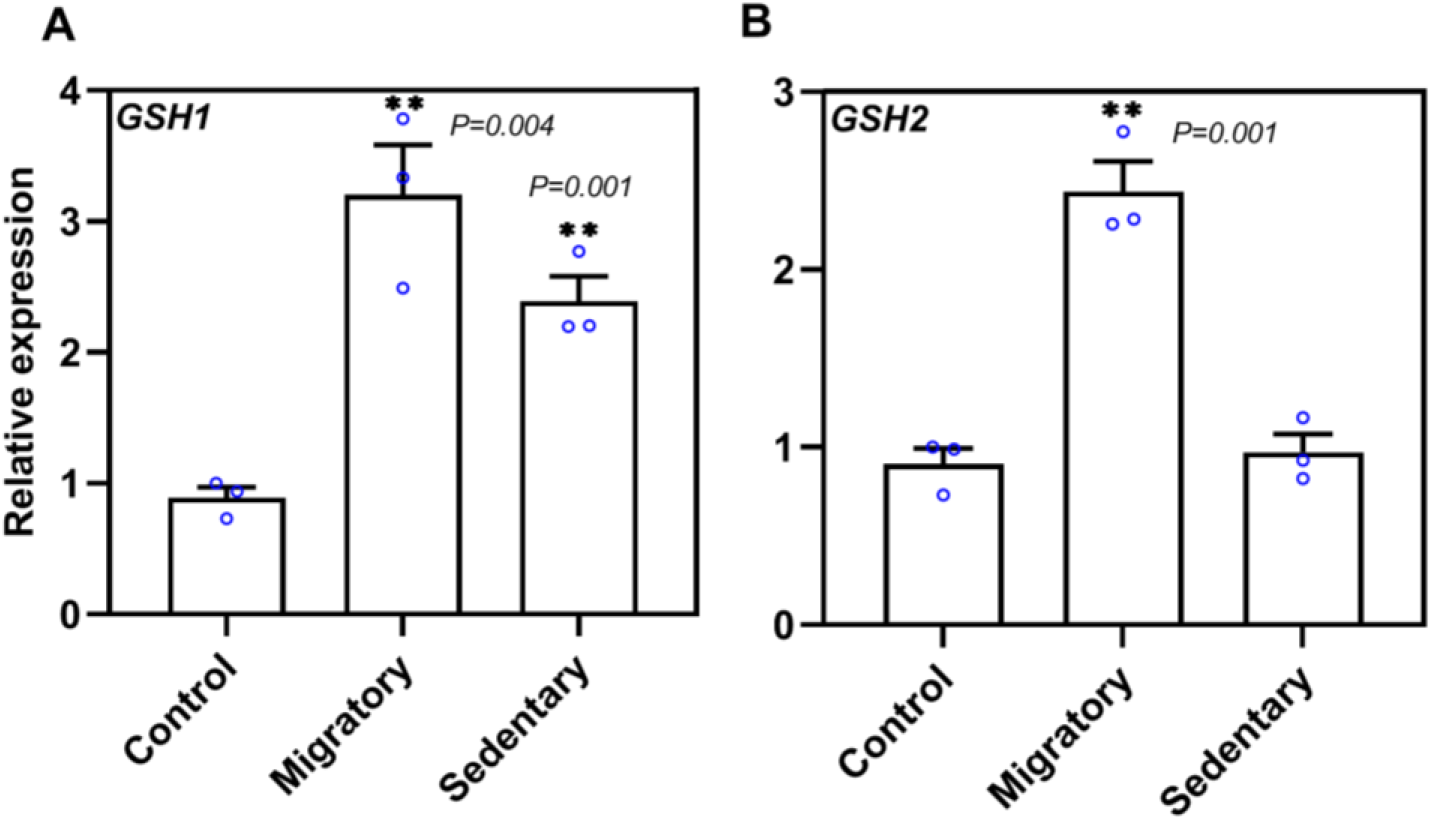
Glutathione (GSH) biosynthesis genes are induced by infection with *Heterodera schachtii*. Validation of changes in expression of genes coding for glutathione biosynthesis enzymes (GSH1 (A) and GSH2 (B) during the migratory and sedentary stages of nematode infection via RT-qPCR. Values represent relative expression levels upon CN infection, with the value of 1^st^ replicate in uninfected control roots (Col-0) set to 1. The mRNA levels were measured in three technical replicates per sample. The transcript level of each gene was normalized to that of the Arabidopsis housekeeping gene *18S*. Data are presented as the mean ± SE for three independent experiments. Each biological replicate (blue circles) contained a pool of hundreds of small root segments with infection sites at 10 hpi (‘Migratory’) or syncytia at 10 dpi (‘Sedentary’). Data were analyzed with a two-tailed Student’s *t*-test (*P* < 0.05), and asterisks represent significant differences compared to the uninfected control root.

### Altered glutathione levels in plants influence cyst nematode infection and development

To assess whether glutathione levels in Arabidopsis affect CN parasitism, we screened an allelic series of *GSH1* mutants in nematode infection assays by measuring multiple nematode susceptibility parameters. We grew plants under sterile conditions in agar medium, and when the roots spread through the agar, we inoculated them with 60–70 J2 nematodes per plant. At 14 dpi, we counted the average number of females, a widely accepted parameter for nematode susceptibility under *in vitro* conditions. The average number of females per centimeter of root length was significantly higher in *zir1* and *cad2* than in the Col-0 control (**Fig. 2A**). However, *rax1* and *pad2* plants showed no difference in the number of females as compared to Col-0 (**Fig. 2A**). Next, we measured the size of female nematodes and female-associated syncytia at 14 dpi. There was no significant difference in either parameter in any lines carrying allelic mutations in *GSH1*, except for *rax1*, which, surprisingly, displayed a slight decrease in the size of female-associated syncytia relative to the wild-type (**Fig. 2B-C**). Finally, we compared cyst egg contents and cyst size at 42 dpi. We detected substantially more eggs per cyst in *pad2, cad2*, and *zir1* than Col-0, but not in *rax1* plants (**Fig. 2D**). However, the cyst size was unaffected in all *GSH1* mutants examined (**Fig. 2E**). Taken together, these findings suggest that glutathione depletion renders plants more susceptible to CN infection.

**Fig. 2.**
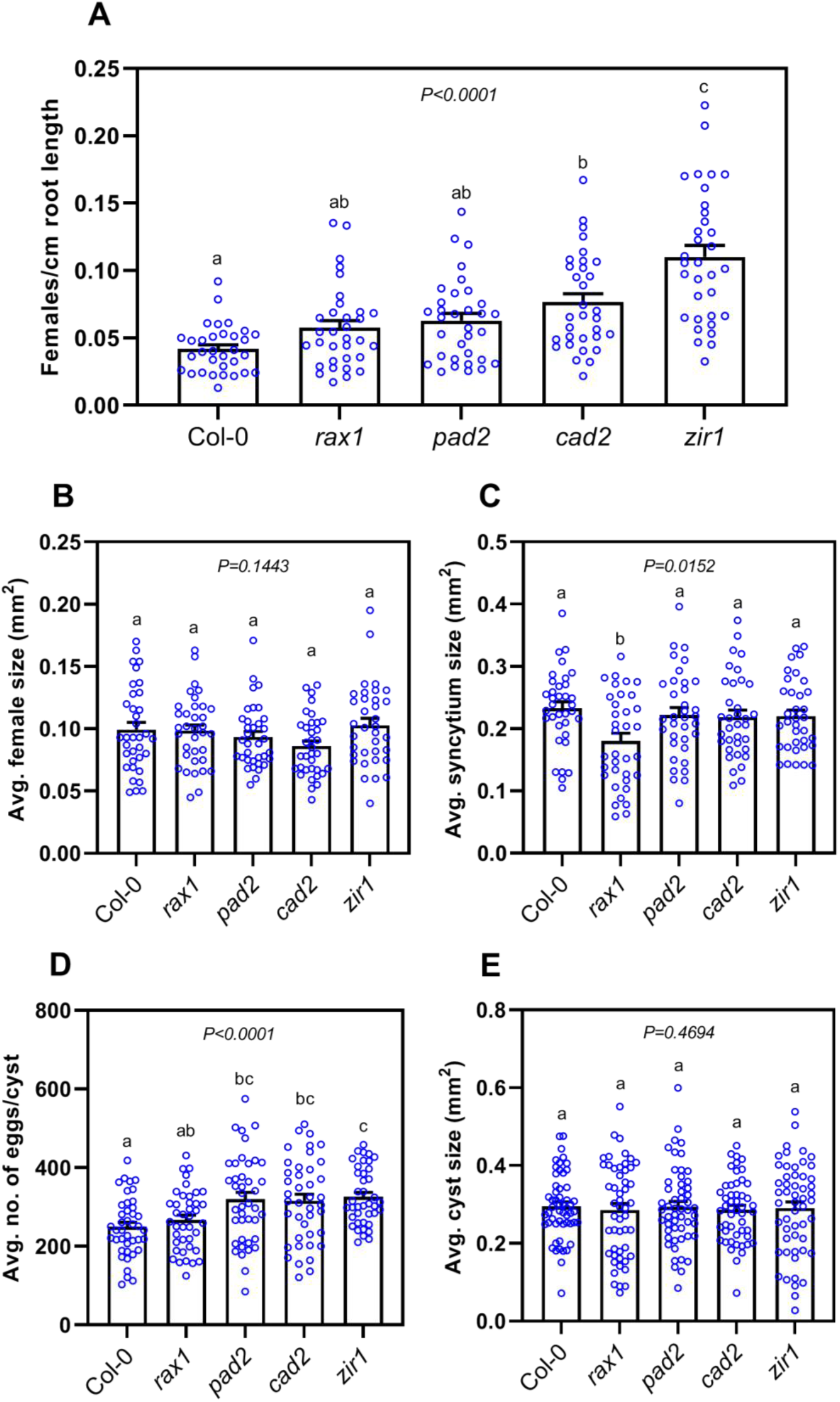
Effect of glutathione (GSH) biosynthesis mutations on the susceptibility of Arabidopsis to *Heterodera schachtii*. (**A**) Nematode infection assay showing number of females/cm root length of *GSH1* mutant lines versus wild-type Col-0. Approximately 60–70 *H. schachtii* infective juveniles were inoculated onto 12-day-old Arabidopsis plants. The number of females per root system was counted at 14 days post-inoculation (dpi), and the infection rate per centimeter of root length was determined after scanning the roots using the WinRhizo root image analysis system. (**B**) Average size of female nematodes at 14 dpi. Approximately 30–40 female nematodes were randomly selected, and their outlines measured for each biological replicate. (**C**) Average size of syncytia at 14 dpi. Approximately 30–40 female-associated syncytia in each replicate were measured using a stereo microscope. (**D**) Average number of eggs per cyst at 42 dpi. For each biological replicate, 40-45 cysts were randomly selected and crushed between slides, their contents transferred to a counting dish, and the number of J2s/eggs counted under a stereomicroscope. (**E**) Average cyst size (50–55 per repetition) at 42 dpi. Bars represent mean ± SE. All data were analyzed using one-way analysis of variance (ANOVA) followed by Tukey’s HSD post-hoc test. Different letters indicate significantly different means (95% confidence). Data represent one of three independent experiments with similar results.

### Changes in the susceptibility of glutathione-deficient mutants are unrelated to CN attraction

To elucidate whether the changes in susceptibility of the *GSH1* mutants to *H. schachtii* were associated with the attractiveness of the plant roots, we performed nematode attraction assays using agar discs containing root exudates from 12-day-old mutants and wild-type control plants. Agar discs containing root exudates of Col-0 plants attracted more J2s than control agar discs (without roots), indicating that nematodes were able to sense the signal from the root exudates (**Supplementary Fig. S1**). However, we observed no significant difference in attractiveness towards the root exudates for any *GSH1* mutant compared with control plants (**Supplementary Fig. S1**). Thus, the changes in the glutathione-deficient mutants’ susceptibility to CN infection are probably not associated with any factor(s) mediating the plants’ attractiveness to nematodes.

### Thiol levels are altered in *GSH1* mutants, regardless of CN infection

Arabidopsis plants harboring mutant alleles of *GSH1* contain constitutively reduced foliar glutathione levels (Parisy et al., 2007; Shanmugam et al., 2012). We explored the extent to which glutathione accumulation is affected in roots due to mutations in *GSH1*. For this purpose, we collected uninfected root tissues from 12-day-old plants and measured glutathione levels. We observed a similar trend in root glutathione levels in *GSH1* mutants as in the foliar levels reported previously (Parisy et al., 2007; Shanmugam et al., 2012), but these levels differed: *rax1* had the highest glutathione level (54% of Col-0), followed by *pad2* (46%), *cad2* (31%), and *zir1* (30%; **Fig. 3A**). Next, we measured glutathione levels in root pieces with infection sites at 10 hpi (migratory stage) and 10 dpi (sedentary stage). Glutathione levels were significantly higher in infected Col-0 at 10 hpi than in uninfected control roots at 10 hpi. However, glutathione returned to uninfected control levels at 10 dpi. In contrast to Col-0, no increase was detected in glutathione levels in the roots of any of the *GSH1* mutant upon nematode infection (**Fig. 3A**).

**Fig. 3.**
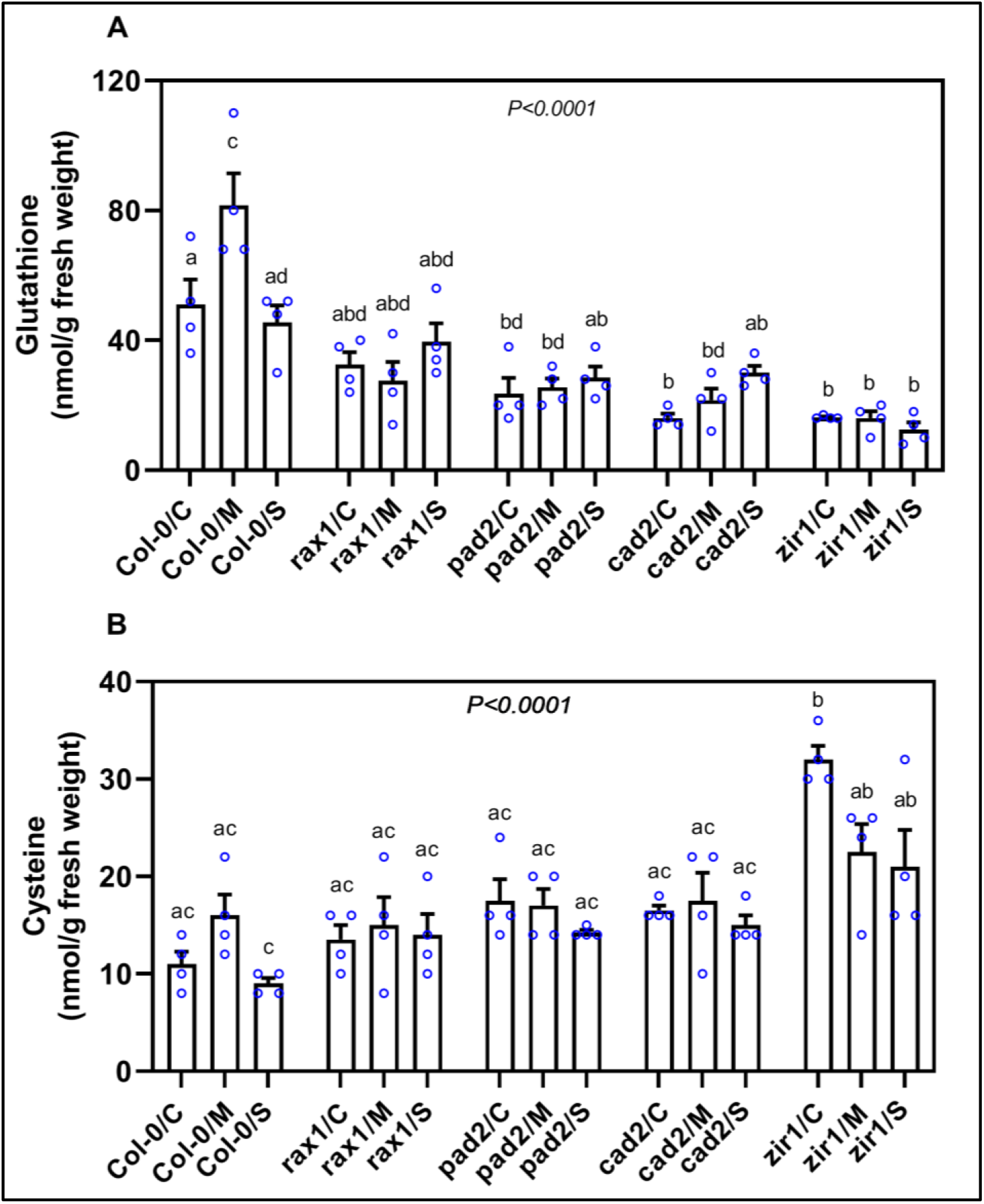
Thiol contents in uninfected roots and cyst nematode infection sites in the *GSH1* mutants *rax1, pad2, cad2*, and *zir1*. Several hundred small root segments (∼0.2 cm) containing infection sites at the migratory (10 hpi) and sedentary (10 dpi) stages were collected for measurement of glutathione (**A**) and cysteine (**B**) contents by HPLC. Data points represent four independent experiments (means ± SE). Different lowercase letters denote significant differences, as determined by ANOVA (*P* < 0.05) followed by Tukey’s HSD post-hoc tests. C, control; M, migratory; S, sedentary.

Cysteine functions as a metabolic precursor for numerous essential biomolecules, such as glutathione, glucosinolates, and phytoalexins (Meyer and Hell, 2005; Rausch and Wachter, 2005). Because the amount of cysteine often inversely correlates with the amount of GSH (Speiser et al., 2018), we measured cysteine levels in roots during different stages of CN infection. Surprisingly, the uninfected roots of *GSH1* mutants contained wild-type levels of cysteine except for *zir1*, which showed a significant increase in cysteine compared with Col-0 (**Fig. 3B**). Similarly, we observed no notable changes in cysteine accumulation in infected root segments of mutant plants at 10 hpi or 10 dpi compared with Col-0 (**Fig. 3B**).

### Glutathione-depleted mutants have impaired basal defense to CN

We hypothesized that the hypersusceptibility of GSH-deficient mutants might be due to impaired expression of genes in defence-related pathways. Therefore, we assessed the transcript abundance of following six plant basal defense marker genes that are strongly up-regulated during the migratory stage of infection (Shah et al., 2017; Mendy et al., 2017): *NPR1*, a salicylic acid signaling gene (Yan and Dong, 2014); *ACS2*, which is involved in ethylene signaling (Yamagami et al., 2003); *JAZ10*, a jasmonic acid signaling gene (Chini et al., 2007); and three genes associated with indole-glucosinolate and camalexin biosynthesis pathways, including the genes encoding GSTF6 [mediates the conjugation between indole-3-acetonitrile (IAN) and GSH (Su et al., 2011)], CYP71B15 [PAD3; catalyzes the conversion of dihydrocamalexic acid (DHCA) to its final form (camalexin) in the camalexin biosynthesis pathway (Schuhegger et al., 2006)], and CYP81F2 [involved in the catabolism of indol-3-yl-methyl glucosinolate (Clay et al., 2009)]. The results from RT-qPCR showed no significant differences in the expression levels of any of the tested marker genes between wild-type uninfected roots and those of the *GSH1* mutants (**Supplementary Fig. S2**). Next, we sampled small root segments (∼0.2 cm) containing infection sites at the migratory stage of CN infection (10 hpi) and analysed gene expression by RT-qPCR. As previously observed (Shah et al., 2017; Mendy et al., 2017), expression of all tested genes was significantly increased in Col-0 upon infection compared with uninfected control roots (**Fig. 4A–F**). In comparison to Col-0, normal increase in transcript levels of all tested genes was impaired in *pad2, cad2*, and *zir1* (**Fig. 4A–F**). Interestingly, *rax1*showed a normal or an even more pronounced increases in defence marker gene expression upon infection (**Fig. 4A)**. Together, these findings suggest that glutathione is involved in activation of plant defense responses, including indole-glucosinolate and camalexin biosynthesis pathways, upon CN infection in Arabidopsis roots.

**Fig. 4.**
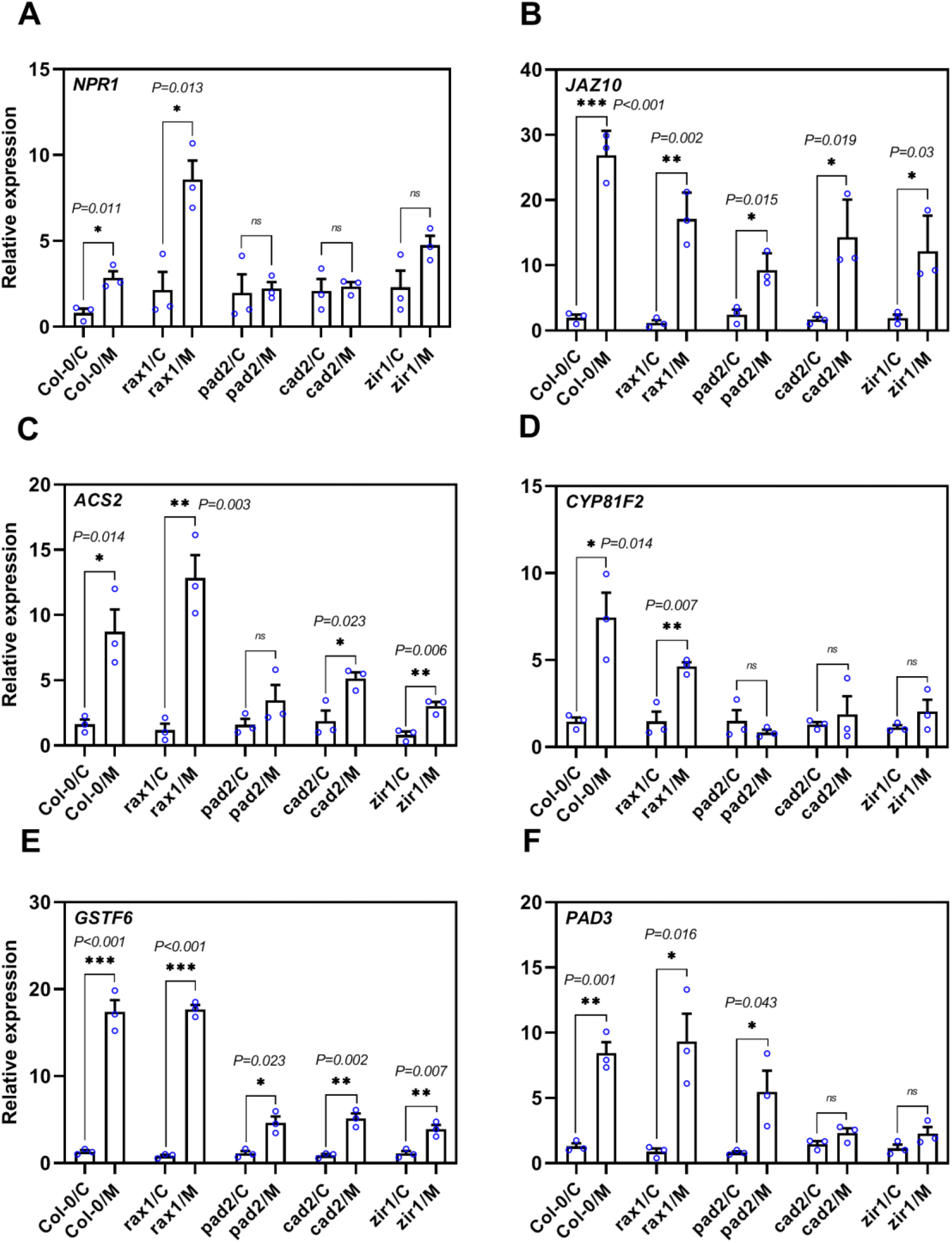
Reduced glutathione levels lead to impaired basal defense responses. **(A-F)** Expression of defense marker genes in response to *Heterodera schachtii* infection during the migratory stage in roots of GSH-deficient mutants. Several hundred root segments (∼0.2 cm) containing infection sites were collected at 10 hpi and the expression levels of *NPR1, JAZ10, ACS2, CYP81F2, GSTF6*, and *PAD3* were analyzed by RT-qPCR. Expression levels of the genes of interest were normalized to the Arabidopsis housekeeping gene *18S rRNA*. Data represent relative expression levels of the indicated genes, with the value for uninfected roots of each genotype set to 1. Asterisks represent significant difference in expression levels from the corresponding uninfected control roots. Data represent means ± SE (*n* = 3 biological replicates, \**P* < 0.05 using two-tailed Student’s *t*-test). C, control; M, migratory stage of infection; ns, non-significant.

### CN parasitism causes shifts in the redox state of glutathione

The glutathione-dependent fluorescent probe Grx1-roGFP2 is commonly used to monitor *E*_GSH_ at the subcellular level *in vivo* (Meyer et al., 2007; Schwarzländer et al., 2008; Müller-Schüssle et al., 2021). To assess whether CN parasitism causes changes in *E*_GSH_, we performed ratiometric analysis of Grx1-roGFP2 (Gutscher et al., 2008; Wagner et al., 2019) expressed in the cytosol via confocal microscopy following excitation at 405 and 488 nm. For calibration, we exposed uninfected roots to reducing (10 mM DTT) and oxidizing (5 mM DPS) reagents. We found that the 405/488 nm fluorescence ratio in Col-0 roots was slightly increased at 10 hpi indicating a less negative *E*_GSH_ (**Fig. 5A**). Together with the increase in total GSH during the migratory state (**Fig. 3A**) this result indicates a slight oxidation of the glutathione pool (**Fig. 5A**). A similar trend albeit with larger variance towards an even more pronounced oxidation was found in of *rax1* (**Fig. 5A)**. In both *pad2* and *cad2* under non-infected control conditions, an increased ratio indicated a less reducing *E*_GSH_, which was expected from the pronounced decrease in total glutathione (Meyer et al., 2007). While *cad2*, which also displayed enhanced susceptibility to *H. schachtii*, showed no change in Grx1-roGFP2 oxidation upon infection, *pad2* surprisingly showed a more reducing *E*_GSH_ after infection (**Fig. 5A**).

**Fig. 5.**
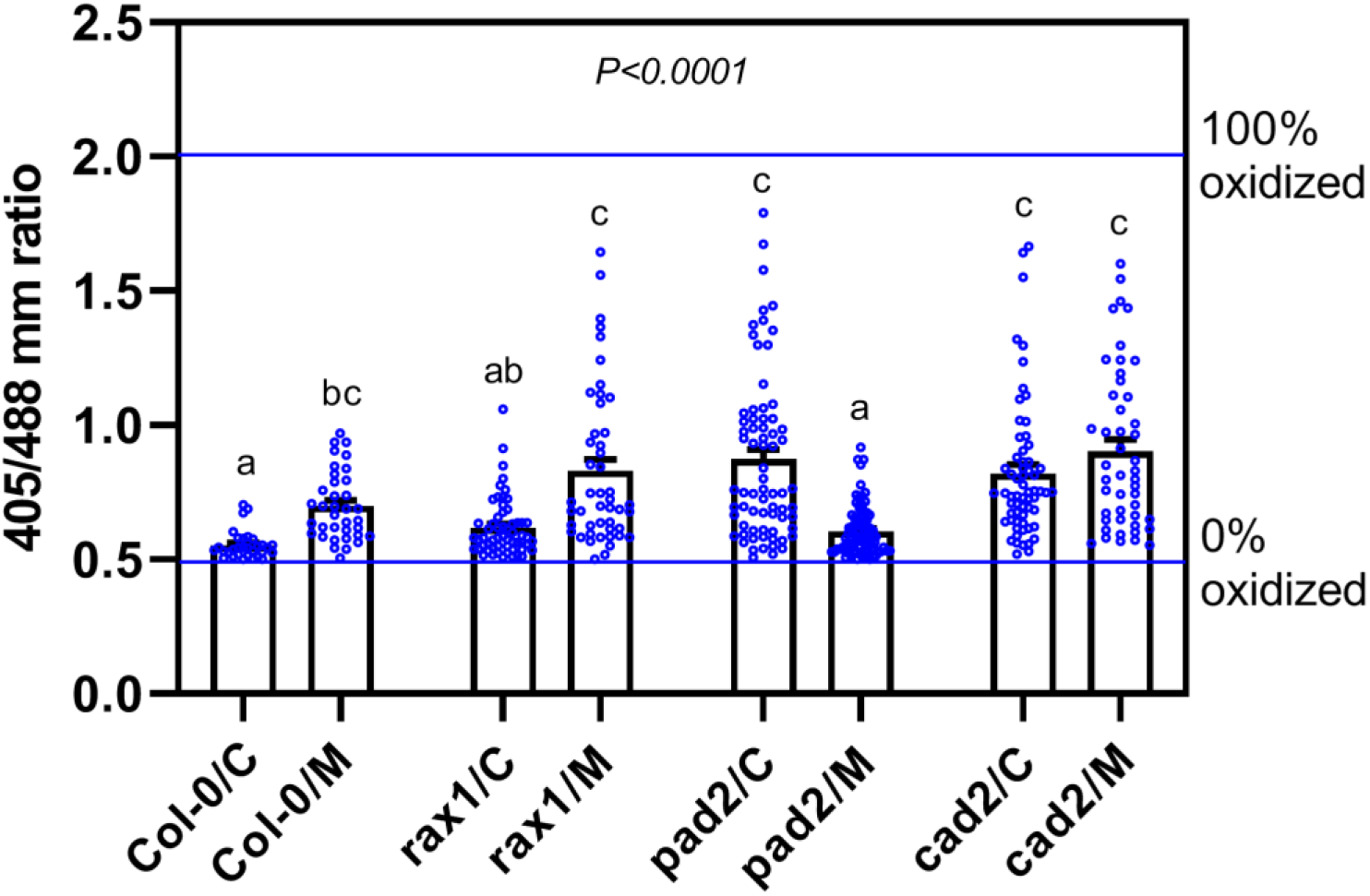
Response of the glutathione redox potential in the cytosol of uninfected and infected (10 hpi) measured with Grx1-roGFP2. The fluorescent probe Grx1-roGFP2 was subsequently excited at 405 and 488 nm and fluorescence recorded at 505-530 nm. For calibration (determining 0% oxidation and 100% oxidation of Grx1-roGFP2), 5 mM DPS solution was used as an oxidizing agent, and 10 mM DTT solution was used as a reducing agent. The respective ratios are depicted as horizontal lines in the diagram. Data points denote individual ratio measurements from regions of interest near infection sites. Results are from three biological replicates (bar chart depicts means ± SE). Different lowercase letters denote significant differences, as determined by one-way ANOVA (*P* < 0.05) followed by Tukey’s HSD post-hoc tests. c, Control; M, migratory.

### GSH-mediated camalexin levels in roots determine plant susceptibility to CN infection

To explore whether there is a link between glutathione and nematode-induced camalexin biosynthesis, we used HPLC to measure camalexin levels in root segments at 10 hpi and in uninfected roots. In uninfected roots, camalexin levels were low and no significant differences in camalexin content were detected between Col-0 and glutathione-deficient mutants. However, we observed an almost 100-fold increase in camalexin content upon CN infection in Col-0 (**Fig. 6)**. Interestingly, an even more pronounced increase was found in camalexin level in *rax1* roots upon CN infection (**Fig. 6**). However, in both *cad2* and *zir1* roots, the increase in camalexin levels was significantly lower than for Col-0 following CN infection, and in *pad2* roots, induction was the same (**Fig. 6**).

**Fig. 6.**
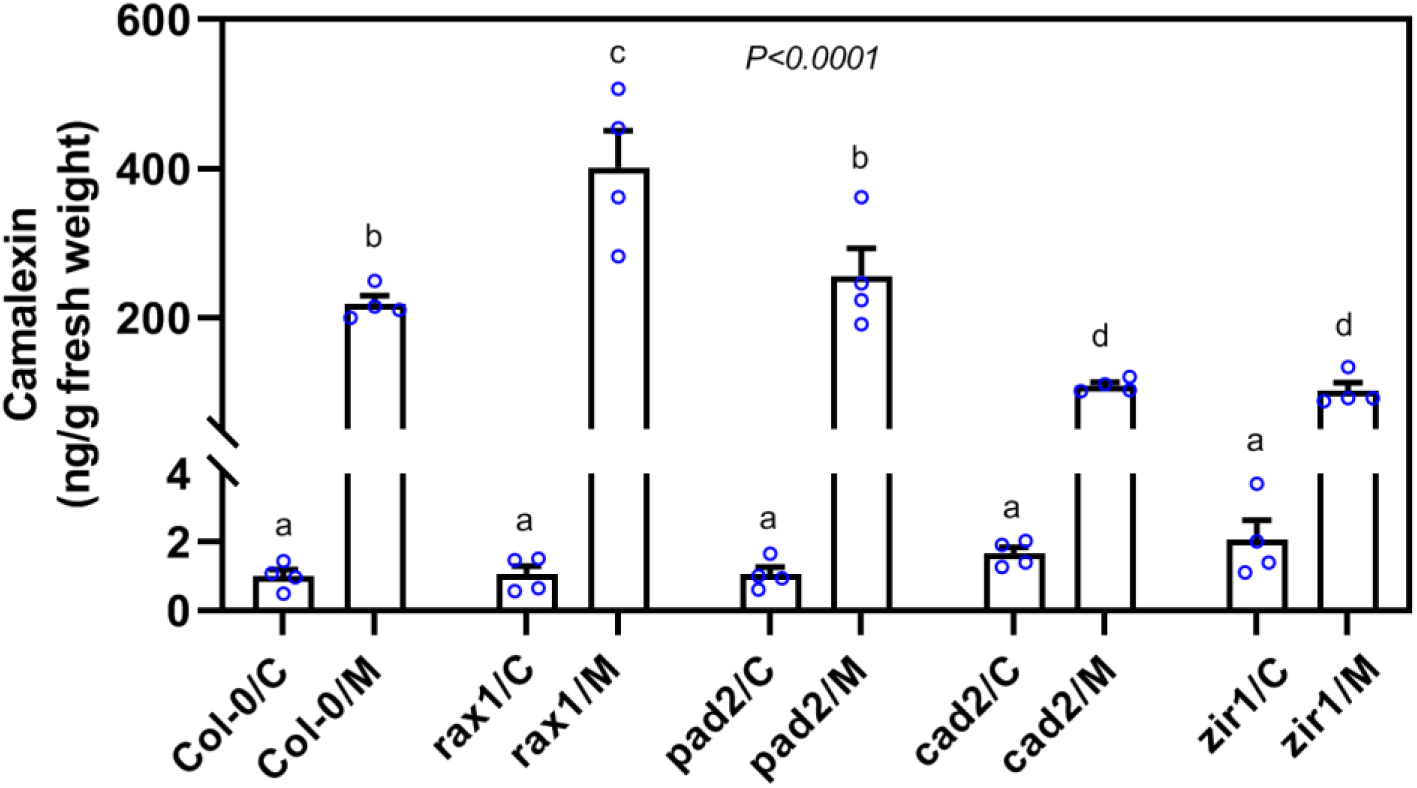
Camalexin levels in uninfected and cyst-nematode-infected roots during the migratory stage (10 hpi) in GSH-deficient mutants measured by HPLC. Data points represent four independent experiments (means ± SE). Data were analyzed by one-way analysis of variance (ANOVA) followed by Tukey’s HSD post-hoc multiple comparisons test (*P* < 0.0001). Similar lowercase letters denote no significant differences. C, Control; M, migratory stage of infection.

## Discussion

In the present work, we utilize Arabidopsis GSH biosynthesis mutants to examine how glutathione contributes to plant susceptibility to CN infection. We determined that the expression of *GSH1* and *GSH2* is significantly increased in response to CN infection during the migratory stage of CN infection. CNs cause extensive damage to host root tissues during their migration inside the roots (Shah et al., 2017). In turn, ROS production and the detoxification machinery are simultaneously switched on to fine-tune the activation of a network of plant defense responses (Molinari and Miacola, 1997; Ithal et al., 2007; Mazarei et al., 2011; Siddique et al., 2014). Increased expression of *GSH1* and *GSH2* is therefore may be a response to the high demand for glutathione to re-establish ROS homeostasis after an infection or for synthesis of the sulfur-containing phytoalexin camalexin during the migratory stage of infection (Parisy et al., 2007; Siddique et al., 2014).

While null mutants of *GSH1* in Arabidopsis are embryo-lethal (Cairns et al., 2006), five distinct mutant alleles (*rml1, rax1, pad2, cad2*, and *zir1*) show a partial decrease in glutathione production (Glazebrook and Ausubel, 1994; Cobbett et al., 1998; Ball et al., 2004; Vernoux et al., 2000; Shanmugam et al., 2012; Bangash et al., 2019). Of these, *rml1* contained <5% of the wild-type levels of glutathione and showed aborted root growth (Vernoux et al., 2000). Because this severe phenotype compromised the analysis of *rml1*, we analyzed plant–CN interactions in the four other *GSH1* mutants (*rax1, pad2, cad2*, and *zir1*). We found that glutathione content increased significantly during the migratory stage of infection in Col-0 compared with uninfected control plants, which supports our observation of increased GSH biosynthetic gene expression. However, glutathione levels were consistently low in *GSH1* mutant plants *(rax1, pad2, cad2*, and *zir1*) with and without infection. Notably, three out of four *GSH1* mutants *(pad2, cad2*, and *zir1*) also showed enhanced susceptibility to CN, suggesting a positive role for glutathione in host defense activation upon CN infection.

Cellular redox homeostasis relies on the equilibrium between the oxidized and reduced forms of glutathione and influences many cellular processes via direct or indirect regulation at the gene or protein level (Cobbett et al., 1998; Mou et al., 2003; Ball et al., 2004; Jez et al., 2004). The increased glutathione accumulation in response to infection suggests that the hypersusceptibility of the glutathione mutants to CN might be associated with disturbances in the cellular redox status of the host plant. Monitoring the fluorescent *E*_GSH_ sensor Grx1-roGFP2 *in vivo*, we found a more oxidized probe during the migratory stage of nematode infection in Col-0. Interestingly, *rax1* mutants showed an oxidative shift of *E*_GSH_, as Col-0, and partially retained induction of defense gene expression. In contrast, this oxidative *E*_GSH_ shift at 10 hpi was absent in the more susceptible *cad2* and *pad2* plants. This finding suggests that glutathione deficiency interferes with pathogen-triggered signaling events that are crucial for successful nematode parasitism, leading to increased plant susceptibility to CN.

The importance of GSH for camalexin accumulation and for disease resistance has been previously demonstrated (Parisy et al. 2007; Glazebrook and Ausubel, 2007). Further, loss-of-function camalexin biosynthesis mutants have been shown to display enhanced susceptibility to CN (Ali et al., 2014; Shah et al., 2017). Here, we found that the up-regulation of two key camalexin biosynthesis genes in response to CN infection was impaired in GSH-deficient mutants as compared with Col-0 control plants. Further, camalexin accumulation in response to CN infection was significantly lower in the roots of in *cad2* and *zir1*. Based on these data, we propose that an adequate level of GSH is required for camalexin accumulation, which in turn play a role in the plant’s defense against CN infection.

Surprisingly, the *rax1* mutant showed a different biochemical phenotype than the other *GSH1* mutants, with significantly higher levels of camalexin during early nematode parasitism. Moreover, the cellular redox state was more oxidized in *rax1* than in the other mutants, pointing to oxidative stress conditions, which might ultimately influence camalexin accumulation in these plants. In our experiments, *rax1* was noted to contain 54% of the wild-type level of glutathione, the highest amount produced by any *GSH1* mutant, indicating that it is able to utilize cysteine to maintain glutathione biosynthesis at a level sufficient to resist nematode invasion. Indeed, *rax1* showed a significant reduction in the size of female-associated syncytia. Moreover, *rax1* shows increased *ASCORBATE PEROXIDASE2* (*APX2*) expression in response to wounding (Ball et al., 2004). Loss-of-function *apx2* mutants are compromised in ROS production during oxidative stress (Suzuki et al., 2013). *APX2* expression is probably triggered by *H. schachtii* during its destructive invasion and migration, leading to excessive ROS accumulation in the roots of *rax1*. Although the *rax1* mutant displays lower ROS accumulation under non-stress conditions, and CNs can utilize ROS for successful parasitism, the mutant’s overproduction of ROS upon nematode infection (caused by the misregulation of the ascorbate– glutathione cycle, resulting in oxidative stress and enhanced camalexin production), might hamper nematode development (Noctor and Foyer, 1998; Zhao et al., 1998; Tierens et al., 2002; Ball et al., 2004; Siddique et al., 2014).

In summary, we demonstrated that glutathione contributes to the establishment and signaling of plant responses to CNs. Considering our observations, we propose that glutathione depletion is positively correlated with camalexin accumulation in roots upon nematode infection, which perturbs the cellular redox state and suppresses early signaling events that positively influence plant susceptibility to CNs. Glutathione deficiency might also accelerate these processes in favor of the host plant to suppress nematode development. Our findings suggest that the precise regulation of glutathione homeostasis is crucial for mounting an effective plant defense response to nematode infection.

## Materials and Methods

### Plant material and growth conditions

*Arabidopsis thaliana* seeds were disinfected by washing in 2% sodium hypochlorite (w/v) for three minutes, followed by washing with 70% (v/v) ethanol for 5 minutes and rinsing three times consecutively with sterile water. After being dried on sterile Whatman filter paper for 2– 4 h, seeds were stored at 4°C before plating. Sterilized seeds were sown in Petri dishes with agar medium enriched with modified Knop’s nutrient solutions as previously described (Sijmons et al., 1991). Plants were grown under long-day conditions with 16 h of light and 8 h of darkness in a growth chamber at 23°C for CN infection (Siddique et al., 2015).

### Validation of mutant lines

The mutant lines *pad2, cad2, rax1*, and *zir1* used in our study were validated by PCR. The area surrounding the predicted point mutation was amplified by PCR using primers listed in **Supplementary Table S3** and, subsequently, the single point mutation/deletion was verified by sequencing. Genotyping results of the mutants are presented in **Supplementary Fig. S3**.

### Nematode infection assay

*H. schachtii* cysts were harvested from mono-culture on mustard (*Sinapis alba* cv Albatros) roots growing on Knop medium (0.2% w/v). The hatching of the juveniles was stimulated by adding 3 mM ZnCl_2_. Upon three consecutive washes with sterile water, 60–70 *H. schachtii* second-stage juveniles (J2s) were inoculated onto the Knop medium plate containing 12-day-old Arabidopsis plants under sterile conditions. Two plants were used in one Petri dish and experiments were repeated at least three times independently, with 20–30 plants per genotype in each replicate. The numbers of female nematodes per plant were counted using a stereomicroscope (Leica Microsystems) at 14 days post-inoculation (dpi). Subsequently, the infection rate per centimeter root length was determined after scanning roots with the WinRhizo root image analysis system. The female nematodes and female-associated syncytia were outlined, and the area was calculated at 14 dpi using a Leica M 165C stereomicroscope equipped with Leica LASv4.3 image analysis software (Leica Microsystems). At 42 dpi, cysts were randomly selected and crushed in between slides. The contents were then transferred into a counting dish and the number of eggs/J2s were counted using a Leica S4E stereomicroscope. Cyst sizes were measured at 42 dpi.

### Nematode attraction assay

Nematode attraction assays were conducted on all glutathione-deficient mutants, according to Dalzell et al. (2011), with some modifications. Uniform circular counting wells of 6 mm in diameter, attached through cylindrical tunnels (2.5 mm depth × 20 mm length) were constructed in a 2% (w/v) water agar plate. The cylindrical channels were created by placing a 20 mm long plastic tube constructed from the handle of a regular inoculation loop onto the agar surface immediately after pouring. With the aid of fine forceps, the tubular plastic was removed once the media had solidified. The wells were cut with a small transfer glass pipette at either side of the central channels to create a 6 mm diameter. Agar plugs were excised from the plate close to the roots carrying root exudates from Col-0 and mutant plants raised on Knop medium and subsequently transferred into the counting wells. Around 60–80 J2s were put in the center of the cylindrical linking channel and kept at room temperature in a dark place. Over a 2 h period, the number of nematodes that moved to either one or the other well were counted and considered as attracted by the exudates of the respective agar plug. Experiments were repeated three times independently with six plates each (*n* = 18). The attraction rate (%) was calculated from the total number of nematodes applied.

### Biochemical analysis

For biochemical analysis, 12-day-old Arabidopsis plants were inoculated with H. *schachtii*. Tiny root segments containing infection sites around the nematode at 10 hpi and female-associated syncytia at 10 dpi were dissected under a stereomicroscope and collected in liquid N2. The root tip or lateral root primordia were excluded during collection. Similarly, the control sample was collected from the corresponding root segments of uninfected plants. Cysteine and glutathione in plant tissues were extracted and quantified by HPLC, according to Anoman et al. (2019). Similarly, samples were collected at 10 hpi, and camalexin was measured as described previously (Koprivova et al., 2019).

### Reverse transcription quantitative PCR

To collect root tissues for gene expression analysis, Arabidopsis Col-0 plants were raised and inoculated using 4-day-old J2s of *H. schachtii*, as described earlier. Several hundred tiny root pieces (∼0.2 cm) containing infection sites around the nematodes at 10 hpi were dissected under a stereomicroscope and collected in liquid N_2_. Total RNA was extracted from the frozen root tissues using an RNeasy Plant Mini kit (Qiagen, Germany) as per the manufacturer’s instructions. The RNA concentration was checked with a NanoDrop (Thermo Fisher Scientific), and reverse transcription was performed using a High Capacity cDNA Reverse Transcription Kit (Life Technologies, cat.no. 4368814), as per the manufacturer’s instructions. The quantitative PCR reactions were carried out in a volume of 20 μL comprising 10 μL Fast SYBR Green qPCR Master Mix with uracil-DNA, 6-carboxy-x-rhodamine, and glycosylase (Invitrogen), 0.5 μL of forward primer, 0.5 μL of reverse primer (10 μM), water and 1 μL of cDNA (c. 100 ng), respectively. The reverse transcription quantitative PCR (RT-qPCR) analysis was conducted using a Stepone Plus Real-Time PCR (Applied Biosystems, USA) system based on a two-step amplification protocol, with the following cycling conditions: 95° C for 3 min followed by 40 cycles of 95°C for 10 s and 60° C for 30 s. For each sample run, a melt-curve analysis was done following 95°C for 15 s, 65–95°C with 0.5°C incremental progress yielding a single peak. As a negative control, a water-containing non-template reaction was included. The transcript abundance of targeted genes was computed from three biological replicates per treatment, with three technical replicates for each biological sample. Relative expression was calculated as stated previously (Pfaffl, 2001) by normalizing target gene expression to the abundance of the Arabidopsis housekeeping gene *18S* to calculate fold change. The gene-specific primer pairs used for RT-qPCR are provided in **Supplementary Table 2**.

### Confocal Laser Scanning Microscopy (CLSM) imaging and ratiometric analysis

CLSM imaging and ratiometric analyses in Col-0, *rax1, pad2* and *cad2* root tissues expressing Grx1-roGFP2 were conducted according to Schwarzländer et al. (2008). In brief, the ratio of fluorescence intensity after excitation at 405 nm and 488 nm of the glutathione redox potential sensor Grx1-roGFP2, was examined in uninfected root tissues and nematode infection sites at 10 hpi. The dynamic range of the sensor (405/488nm ratio) was determined by *in situ* calibration (0.5-2.0) and was set as the min/max value of the ratio false color scale. For calibration, 5 mM DPS (2,2’-dipyridyl disulfide) solution as oxidizing and 10 mM DTT (1.4-dithiothreitol) solution as reducing agents were used. Images were captured using a confocal microscope (Zeiss LSM 780) with a 25x-lens using the multitrack mode, with line switching and averaging two frames. The fluorescence of Grx1-roGFP2 was collected between 505–530 nm. RRA v1.2 software (Fricker, 2016) was used for ratiometric image analysis. To minimize impact by cell wall autofluorescence, values were measured on regions of interest (ROIs) in nuclei of cells near infection sites. Values outside the min/max value of the ratio (0.5-2.0) were removed. Statistical analysis was performed on log-transformed values.

### Statistical analysis

All data were analyzed using the statistical software GraphPad Prism v.8.4.3 for Windows. Infection assays were carried out with 20–30 plants for each genotype and replicated at least three times independently. The normality of data was assessed using the Kolmogorov–Smirnov test/Shapiro–Wilk test (α <0.05). The specific statistical method used for the data set of each experiment is described in the figure legends. Data are presented as mean ± SE, and corresponding *P* values are indicated either in the figure or in the figure legends.

## Acknowledgements

SS’s research was supported by Deutsche Forschungsgemeinschaft (DFG) (Project number 287570125). We thank Bastian Walter (University of Cologne) for technical assistance with thiol analysis. Research in SK’s lab is funded by the Deutsche Forschungsgemeinschaft (DFG) under Germany’
ss Excellence Strategy (EXC 2048/1) – project 390686111. M. Shamim Hasan was supported by a fellowship from German Academic Exchange Service (DAAD; 91525252).

