## Supplementary material for "Glutathione contributes to plant defense against parasitic cyst nematodes": Supplemetary

| **Table S1:** | Overview of glutathione biosynthesis gene *GSH1* and *GSH2* expression patterns in Arabidopsis roots at migratory and sedentary stage of *H. schachtii* infection in published transcriptomic data. |
| --- | --- |
| **Figure S2:** | Nematode attraction assays towards root exudates of glutathione-deficient mutant plants. |
| **Figure S3:** | Genotyping results of glutathione deficient mutants (*rax1*, *pad2, cad2,* and *zir1*) used in this study. |
| **Table S2:** | List of primers used for validation of the mutant lines tested in this study. |

**Table S1. Overview of glutathione biosynthesis gene GSH1 and GSH2 expression patterns in Arabidopsis roots at migratory and sedentary stage of *H*. *schachtii* infection in published transcriptomic data.**

| **Gene** | **Locus** | **Fold change compared with uninfected control** | |
| --- | --- | --- | --- |
|  |  | **Migratory (10 hpi)** | **Sedentary (5+15 dpi)** |
| *GSH1* | At4g23100 | 2.1* | 6.5* |
| *GSH2* | At5g27380 | 2.1* | 0.8 |

For the migratory stage, root sections containing infection sites at 10 hours post infection (hpi) were analyzed and compared with uninfected control roots (Mendy et al., 2017). For the sedentary stage, microaspirated syncytia at 5 and 15 days post-infection (dpi) were pooled and compared with control roots (Szakasits et al., 2009). Asterisks indicate significant difference to control.

**Fig. S1. Nematode attraction assays towards root exudates of glutathione-deficient mutant plants.** Nematode attractiveness to root exudates of *GSH1* mutants compared with Col-0 plants (control = control agar). Experiments were repeated three times independently for each mutant with six plates each (*n* = 18). The attraction rate (%) was calculated from the total number of applied nematodes. Bars represent mean ± SE. Data were analyzed using two-tailed Student’s t‐test (P < 0.05). Asterisks indicate significant differences (*P* < 0.05).

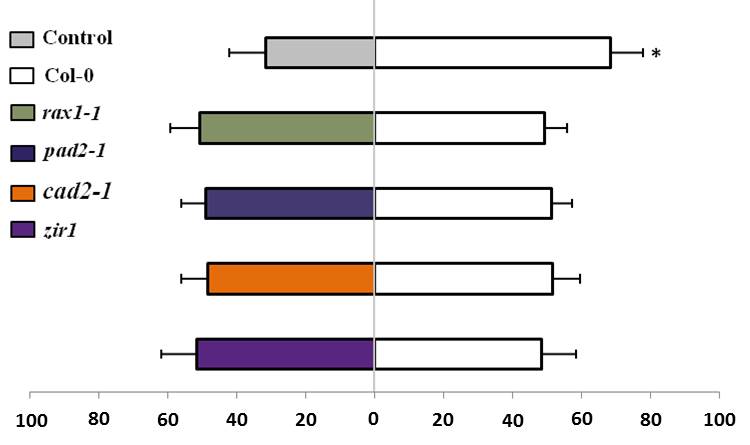

**Fig. S2. Reduced glutathione levels do not impair basal defense responses in uninfected roots.** Expression of defense marker genes in uninfected roots of GSH-deficient mutants.

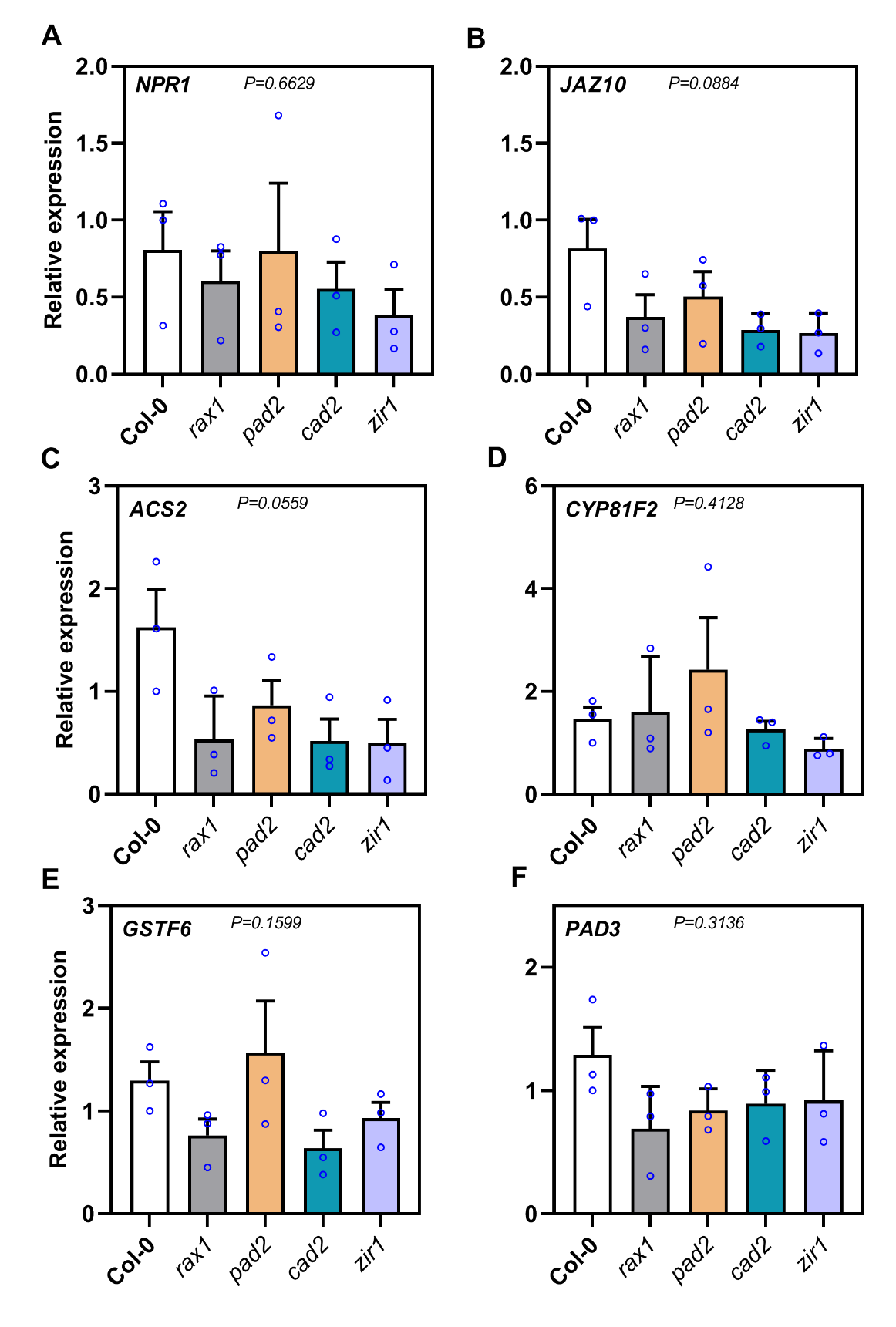

**Fig. S3. Genotyping results of glutathione deficient mutants (*rax1*, *pad2, cad2* and *zir1*) used in this study.** The GSH1 mutants, *rax1*, *pad2,* and *zir1* have single point mutations leading to changes in amino acid (AA) synthesis, and the line *cad2* has a 6 bp long deletion. The point or deletion mutations were identified (shown in a red box) via PCR using primers listed in **Supplementary Table S2** and subsequently, the single point mutation/deletion was verified by sequencing. The shown sequences are only for the PCR products amplified by the respective primers.

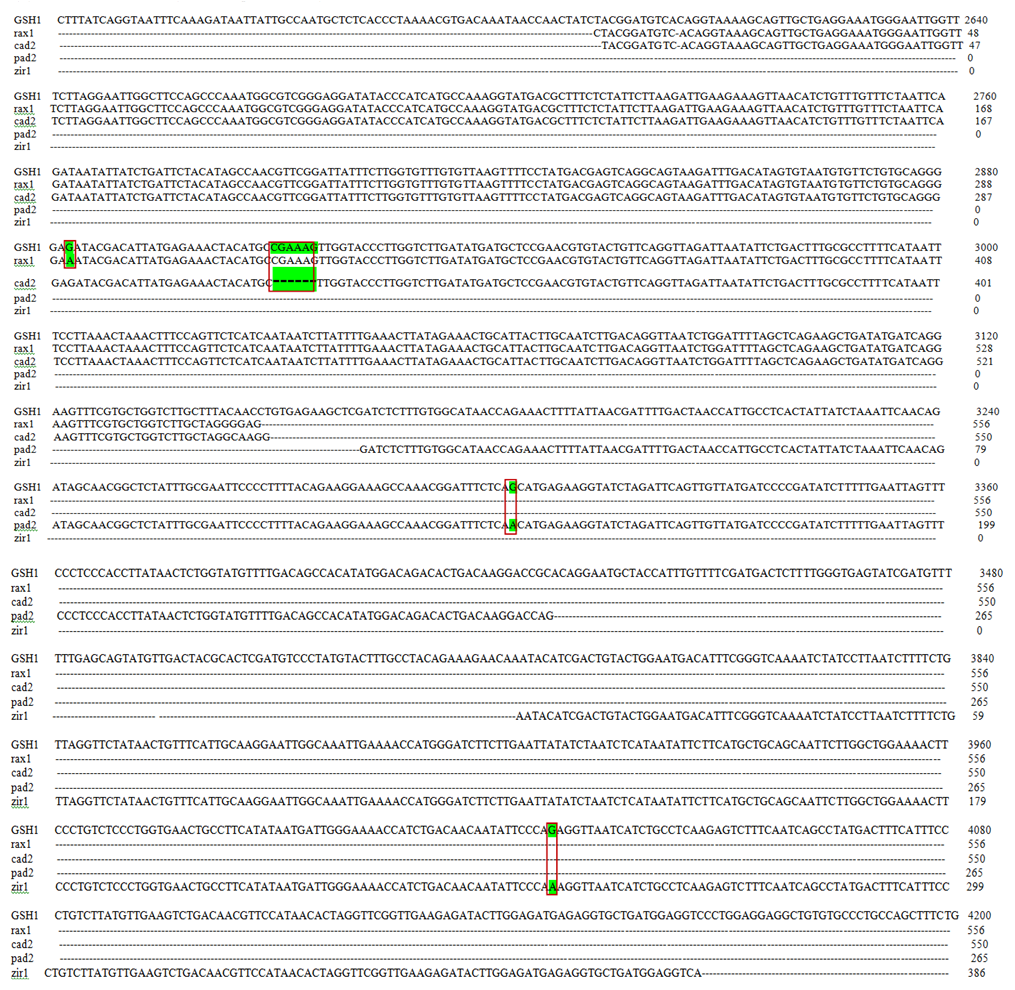

**Table S2. List of primers used in this study.**

| **Gene** | **Locus** | **Forward primer** | **Reverse primer** |
| --- | --- | --- | --- |
| RAX1 | AT4G23100 | TGCTCTCACCCTAAAACGTG | AGCAAGACCAGCACGAAACT |
| PAD2 | AT4G23100 | TCGTGCTGGTCTTGCTTTAC | CGGTCCTTGTCAGTGTCTGT |
| CAD2 | AT4G23100 | TGCTCTCACCCTAAAACGTG | AGCAAGACCAGCACGAAACT |
| ZIR1 | AT4G23100 | ACGCACTCGATGTCCCTATG | GACCTCCATCAGCACCTCTC |
| 18S | 18S RRNA | GGTGGTAACGGGTGACGGAGAAT | CGCCGACCGAAGGGACAAGCCGA |
| NPR1 | AT1G64280 | GAGTTGCACTTGCTCAACGTC | GCTATCTTTACACCCGGTGATG |
| JAZ10 | AT5G13220 | TCGCAAGGAGAAAGTCACTGCAAC | CGATTTAGCAACGACGAAGAAGGC |
| ACS2 | AT1G01480 | GGATGGTTTAGGATTTGCTTTG | GCACTCTTGTTCTGGATTACCTG |
| CYP81F2 | AT5G57220 | ATCGTGCTAGTGAACGCTTG | TTCGTCCGTTACCAAACACC |
| GSTF6 | AT1G02930 | CCAGCCTTTGAAGATGGAGA | CTTGCCAGTTGAGAGAAGGTTG |
| PAD3 | AT3G26830 | GGGTACCATACTTGTTGAGATGG | TTGATGATCTCTTTGGCTTCC |
| GSH1 | AT4G23100 | CCAGCTTTCTGGGTGGGTTT | GCTTCCTTGTAGCCTCTGCG |
| GSH2 | AT5G27380 | TGGCTAAAGCTTGGTTGGAGT | AACCACTGCGACTGCTTGG |
